# MOSAIC: Learning Graph Node Embeddings from Spectrally Isolated Dominant Subspaces of Accumulated Diffusion Operators

**DOI:** 10.64898/2026.09.28.754397

**Authors:** Taha Mohseni Ahooyi, Benjamin J. Stear, Yuanchao Zhang, Aditya Lahiri, James Terry, Shiping Zhang, Alan Simmons, Ryan Corbett, Patricia J. Sullivan, Asif Chinwalla, Christopher Nemarich, Jo Lynne Rokita, Sharon Diskin, Jonathan C. Silverstein, Deanne M. Taylor

## Abstract

Graph embedding transforms complex networks into low-dimensional representations for analysis and downstream machine learning. Existing methods often rely on eigenvalue ordering, random-walk sampling, or neural optimization. We introduce MOSAIC, Multistep Operator Spectral Accumulation with Isolated Components, an unsupervised framework that constructs an accumulated diffusion operator from successive powers of a graph transition matrix. MOSAIC uses Krylov subspace analysis to identify dominant spectral components according to their convergence behavior. This approach emphasizes persistent and spectrally separated diffusion modes rather than selecting embedding coordinates solely by eigenvalue magnitude.

We evaluated MOSAIC using a biomedical graph projection derived from the Data Distillery Knowledge Graph that contained 5,171 human genes and 10,147 phenotypes. Across embedding dimensions from 16 to 128, MOSAIC achieved node-type classification AUROCs from 0.9863 to 0.9926 and exceeded FastRP, Node2Vec, HashGNN, and GraphSAGE. For gene-phenotype link prediction, validation and test associations were withheld before embedding generation. Under progressive removal of all edges or specific edge classes, MOSAIC retained more node-label information than FastRP, including after extreme graph sparsification. In a focused DICER1 analysis, 106 of 137 documented phenotype associations received scores above 0.90, demonstrating concordance with known pleiotropic biology and prioritizing additional relationships for independent evaluation. These results support MOSAIC as a fully unsupervised framework for learning informative graph representations. Although demonstrated using a biomedical graph projection, MOSAIC is domain independent and can be applied to other complex networks.

## Introduction

Knowledge graph (KG) embedding has become a fundamental task in graph representation learning, enabling complex relational structures to be encoded as low-dimensional vector spaces suitable for visualization, clustering, recommendation systems, link prediction, and downstream machine learning applications. The principal objective of graph embedding is to preserve the intrinsic structural properties of a graph while substantially reducing its dimensionality, thereby providing representations that facilitate both computational efficiency and semantic interpretability. Over the past decade, numerous node embedding techniques have been proposed, each emphasizing different aspects of graph topology and information propagation.

To evaluate MOSAIC in a biologically meaningful setting, we applied it to a graph projection derived from the Data Distillery Knowledge Graph. The evaluation network contained 5,171 human genes and 10,147 human phenotypes connected through gene-gene, phenotype-phenotype, and gene-phenotype relationships. This heterogeneous structure supports several complementary tests of graph representation quality, including node-type classification, robustness to progressive loss of different edge classes, and recovery of held-out gene-phenotype associations. We use this network as the primary biomedical demonstration of MOSAIC while retaining a formulation that is applicable to graph representation problems across other scientific and technical domains.

Existing graph embedding methods can broadly be categorized according to the structural information they preserve. Context-preserving approaches, including Node2Vec [1], DeepWalk [2], and Graph Diffusion Embedding Networks (GDENs) [3], exploit random walks or diffusion processes to encode local neighborhoods and higher-order connectivity into continuous vector representations. In contrast, geometry- and manifold-preserving methods such as Laplacian Eigenmaps [4], Graph UMAP [5], and SpectralNet [6] seek embeddings that preserve graph geometry, pairwise distances, and the intrinsic manifold structure by leveraging spectral graph theory or nonlinear dimensionality reduction techniques. More recently, graph embedding research has evolved toward unified frameworks capable of simultaneously preserving contextual, geometric, and manifold information. Representative examples include Hyperbolic Graph Embeddings (HGE) [7], which exploit hyperbolic geometry for hierarchical graph representations, Mixture of Adaptive Graph Experts (MAGE) [8], and GraphMoRE [9], which integrates multi-hop attention with ensemble regularization to capture structural information across multiple scales.

Despite these significant advances, several limitations remain. Many state-of-the-art node embedding techniques require computationally demanding optimization procedures, including full eigendecomposition, iterative Riemannian optimization, or deep graph neural network training, which scale poorly for large graphs [10]. Random walk-based methods frequently depend on carefully tuned hyperparameters, such as walk length, context window size, neighborhood sampling strategy, or manifold initialization, making their performance highly dataset-dependent and reducing reproducibility. Deep graph representation models often sacrifice interpretability for predictive performance, producing embeddings whose geometric relationships are difficult to explain from the underlying graph topology [11,12]. Furthermore, repeated message passing or multi-hop diffusion may lead to over-smoothing, whereby node representations progressively lose their discriminative power as they converge toward similar feature vectors [13]. Many recent embedding frameworks additionally rely on supervised objectives, contrastive learning, or pre-training strategies, limiting their applicability to fully unsupervised exploratory graph analysis [14].

A common characteristic shared by most spectral node embedding methods is that the embedding basis is selected according to the ordering of eigenvalues associated with graph-derived operators, such as adjacency matrices, graph Laplacians, or normalized diffusion operators. This selection criterion originates from classical spectral graph theory, where extremal eigenvalues describe asymptotic properties of the graph, including connectivity, diffusion rates, and graph partitioning. While mathematically well established, eigenvalue ordering alone does not necessarily identify the invariant subspaces that contribute most significantly to the finite-dimensional action of the graph operator on data. In practical applications, particularly for large and irregular graphs, eigenvalues often appear in dense spectral clusters, resulting in eigenvectors that may be numerically sensitive, poorly separated, or insufficiently representative of the graph’s persistent structural organization.

This work introduces an alternative graph node embedding framework based on the dominant, spectrally isolated eigenpairs of a graph diffusion operator named MOSAIC (**M**ultistep **O**perator **S**pectral **A**ccumulation with **I**solated **C**omponents). Rather than selecting eigenvectors solely according to eigenvalue magnitude, the proposed approach identifies invariant subspaces that dominate the operator’s action under Krylov subspace iterations. For the multistep diffusion operator, the resulting spectrum naturally amplifies slowly decaying diffusion modes associated with eigenvalues near unity while suppressing transient components. Consequently, the selected eigenvectors represent persistent diffusion structures that govern long-range information flow through the graph. Spectral isolation further ensures numerical robustness by reducing interactions between neighboring eigenspaces, yielding embedding coordinates that are stable under perturbations and better suited for representing coherent graph topology.

Unlike existing spectral node embedding methods that primarily employ Krylov subspace algorithms as efficient numerical solvers for large eigendecomposition problems, the proposed framework uses the convergence behavior of Krylov iterations as the selection criterion for determining the embedding basis itself. To the best of our knowledge, this perspective has not been explicitly investigated in graph embedding literature, where dominant embedding coordinates are almost universally determined through spectral ordering or matrix factorization. By combining diffusion persistence, spectral isolation, and Krylov-based dominance into a unified embedding criterion, the proposed method provides an interpretable, fully unsupervised, and computationally scalable alternative for graph representation learning that remains closely aligned with the underlying graph dynamics.

### General Definitions

#### 1) Graph multistep operator: finite and infinite forms

Let *G* = *G*(*V, E*), |*V*| = *n* be a graph with adjacency matrix 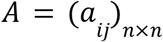, degree matrix *D* = *diag*(*d*_1_,…, *d*_*n*_) and row stochastic transition matrix *P* = *D*^−1^ *A*. The matrix *P* defines a random walk on *G*. The multistep diffusion operator with discount or damping parameter α (where 0 < α < 1) is:

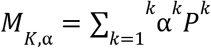

where 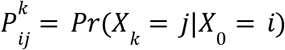 is the probability that a random walk starting at *i* reaches *j* in exactly *k* steps. Therefore

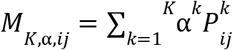

measures the cumulative influence of node *i* on node *j* across all paths of length *K*. Accordingly, *M*_α_ captures local connectivity (*P*), medium-range connectivity (*P*^2^ and *P*^3^) and global connectivity (large *K*s). The infinite multistep diffusion series can be rewritten in closed form as a matrix inverse:

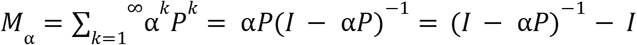

This renders the multistep operator similar to total diffusion (Katz Centrality and the Katz Diffusion Operator) and Personalized PageRank. The (*i, j*)-th entry of this resolvent matrix ((*I −* α*P*)^*−*1^) provides information on how much “diffusion energy” successfully travels from node *i* to node *j* across all possible multistep paths in *G*.

#### 2) Spectral Decomposition of *M*_α_

Assume *P* is diagonalizable i.e. *P* = *U*Λ*U*^*−*1^ where Λ = *diag*(*λ*_1_, …, *λ*_*n*_). Therefore: *P*^*k*^ = *U*Λ^*k*^ *U*^*−*1^and

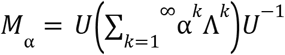

Then

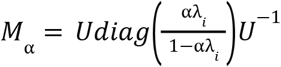

According to these definitions, the diffusion process doesn’t create new directions, therefore the spectral structure of *M*_α_ follows directly from that of *P* and has the same eigenvectors. Decomposition of *M*_α_ in terms of the Eigenvectors of *P* can be rewritten in terms of a function 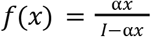

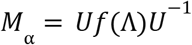

Since *f*(Λ) is diagonal, *M*_α_ and *P* share the same eigenvectors. Given the *r* nontrivial eigenvalues of *P* associated with a connected graph:

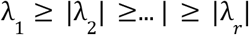

As *f*(*λ*) is monotone increasing in *λ*, the largest eigenvalues receive the largest amplification hence

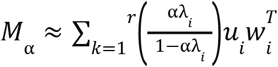

where *r ≪ n*. With *Σ* = *f*(Λ) let

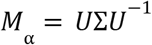

This decomposition allows for an *r*-dimensional vector *x*_*i*_ ∈ *R*^*r*^ associated with nontrivial eigenvalue to represent node *i*-th node of the underlying graph *G* such that 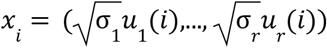 using 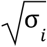 ensures that the squared distance between embedded points optimally approximates the distances in the original high-dimensional space.This diffusion-based embedding is equivalent to embedding with Laplacian with a nonlinear spectral filter 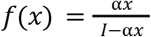:

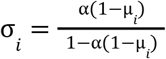

where are µ_*i*_ s are Laplacian eigenvalues.

According the definitions above, a proper embeddings derived from spectral decomposition of *M*_α_ harbors multi-scale information as it incorporates random walks of all lengths. These embeddings provide a level of “community awareness” as vertices from the same cluster would share similar coordinates, so the embeddings preserve the local community geometry. Furthermore, Euclidean distances in the embedding space are hypothesized to approximate diffusion distances (contextual preservation). Additionally the nonlinear spectral filter emphasized slow-mixing structural modes, where the leading eigenvalues capture the best-low ranking approximation of the while small eigenmode attenuation would lead to noise suppression.

#### 3) Constructing the Krylov-Ritz Solver

Suppose we run Arnoldi/Lanczos on *P* (or the multistep version):

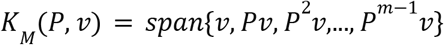

expand

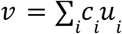

In the eigenbasis.

Then

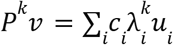

Thus

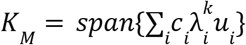

The directions that dominate the Krylov space are those with large 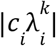 so convergence depends on both:

1. Eigenvalue magnitude
2. Overlap *c*_*i*_ with the starting vector

For a connected graph the leading eigenvector is useless for embedding. It places all nodes at the same coordinate. The second eigenvector is useful. But the third, fourth, fifth, etc. may not necessarily be. A large eigenvalue merely means slow decay under the random walk. It does not necessarily mean discriminative power for node representation.

Suppose

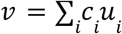

The total energy captured by eigenvector *u*_*i*_ is|*c*_*i*_|^2^

Lanczos/Arnoldi naturally prioritizes modes with large |*c*_*i*_ |^2^ . These are the directions actually excited by the graph signal. In graph signal processing language we can say the Krylov basis adapts to the signal distribution. The pure eigenbasis does not.

As *M*_*K*,α_ = ∑_*k*=1_ ^*k*^ α^*k*^ *P*^*k*^, a Krylov space contains {*v, Pv, P* ^2^ *v*,…, *P*^*m−*1^ *v*} which is exactly the collection of vectors appearing in the diffusion operator, therefore

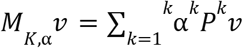

Thus the Krylov space is the natural approximation space for diffusion. This means that the leading Ritz vectors may be approximating the dominant singular directions of the diffusion process rather than the dominant eigenvectors of *P*.

### Hypothesis

For symmetric matrices, Arnoldi convergence satisfies approximately

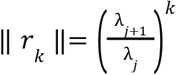

The convergence rate depends on spectral separation. If *λ*_*j*_ is isolated and the Ritz vector converges rapidly. If eigenvalues are clustered then the convergence is generally slow. Thus the eigenvectors discovered first by Arnoldi are often those associated with large spectral gaps. Therefore, fast Krylov convergence is acting as a detector for spectrally isolated graph structures.

In other words, the “best” graph coordinates may correspond to spectrally isolated, dynamically significant modes, rather than simply the largest-magnitude modes. This viewpoint aligns remarkably well with the fact that Krylov-Ritz methods naturally converge first to eigenpairs that are both dynamically important and spectrally distinguishable.

### Dynamic observability

The fastest-converging Ritz vectors correspond not to the largest eigenvalues of eigenvalues of *P*, but to the largest a a diffusion Gramian:

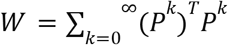

a closely related operator to *M*. Notice that *W* measures total propagated energy across all diffusion times. Its eigenvectors identify the most observable and controllable graph directions. Those directions often produce better embeddings than ordinary spectral coordinates. This viewpoint is closely related to balanced truncation, controllability Gramians, diffusion maps, dynamic mode decomposition and Koopman analysis, and may explain why Ritz convergence correlates with embedding quality. Suppose the Krylov process starts from

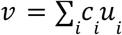

The amount of mode *u*_*i*_ visible to the dynamics is |*c*_*i*_ |^2^ . For symmetric *P*

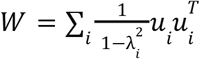

The energy associated with mode *i* is

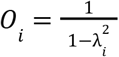

This is the amount of diffusion energy carried by that mode.

### Spectral isolation

We define spectral isolation as

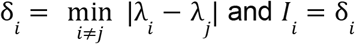

This measures how distinguishable the mode is. Perturbation theory suggest that larger *I*_*i*_ corresponds to a stable eigenvector where smaller *I*_*i*_ would mean there is an ambiguous eigenspace.

### Diffusion persistance

Let

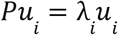

Contribution of mode *u* after *k* diffusion steps is 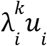

The associated diffusion timescale is

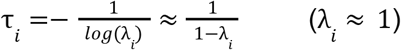

Large τ_*i*_ means slow decay, long-range structure and Cheeger-relevant geometry [15].

We define persistence as:

*P*_*i*_ = τ_*i*_

### Combining observability, isolation and persistence

Let

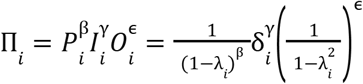

This score increases when

- The mode persists for a long time,
- The mode is spectrally isolated,
- The mode carries large diffusion energy.

This measure explains:

- Cheeger vectors (large 1/(1 *− λ*_*i*_)),
- Krylov-Ritz convergence (large δ_*i*_),
- diffusion-map quality (large diffusion energy),
- robustness under perturbations.

In other words, the graph representation quality through the fastest-converging/most isolated eigenvalues could be rewritten as:

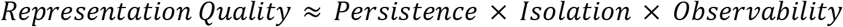

### Example subgraph projection

An example graph consisting of 5,171 human genes and 10,147 human phenotypes (15,318 nodes and 1,577,238 (bidirectional) edges) was created by projecting an induced subgraph with these two node types form our previously published Data Distillery graph (DDKG) [16] in Neo4j graph database management environment [17]. The subgraph was then used to compute node embedding in 16, 32, 64 and 128 dimensions via the FastRP, Node2Vec, HashGNN and GraphSAGE methods utilizing the functions provided by the Graph Data Science (GDS) library plug-in in Neo4j [18]. For MOSAIC trains and prediction, the constructed subgraph was selected to be **undirected**, however **directed** graphs are also fully supported. For MOSAIC computation purposes the subgraph was reconstructed in R (v.4.4.0) [19] using the “igraph” package (v2.1.4) [20].

**Fast Random Projection (FastRP)** node representations were generated using the algorithm implemented in Neo4j Graph Data Science. FastRP was applied to the subgraph projection to produce (16, 32, 64 and 128)-dimensional node embeddings. The algorithm was configured with iteration weights of [0.0, 1.0, 1.0], thereby excluding the initial node representation and assigning equal weight to representations propagated through the first and second graph iterations. The normalization strength and node self-influence were both set to 0.0, eliminating the corresponding normalization and self-representation contributions.

Likewise, **Node2Vec** was applied to the subgraph projection to produce (16, 32, 64 and 128)-dimensional node embeddings. The random-walk procedure used walks of length 80, with 10 walks generated per node, providing the algorithm with repeated samples of graph connectivity. The returnFactor and inOutFactor were both set to 1.0, resulting in an unbiased random-walk strategy with respect to returning to previously visited nodes and exploring outward from the current node. A context window of 10 was used for learning node relationships from the generated walks, while a negative sampling rate of 5 specified five negative samples for each positive training example. The embedding model was trained for one iteration, using an initial learning rate of 0.01 that decayed to a minimum of 0.0001.

Similarly, **HashGNN** was applied to the projected subgraph with heterogeneous graph processing enabled (heterogeneous: true), allowing the embedding procedure to account for multiple node and/or relationship types. The model was run for 20 iterations, with each iteration aggregating information from neighboring nodes. Initial node features were generated using a 32-dimensional feature space with generateFeatures and a density level of 1, while binarizeFeatures transformed the generated features into a 64-dimensional binary representation using a threshold of 0. The final node embeddings were projected into output dimensions of 16, 32, 64 and128, producing multidimensional vectors for each node. The neighborInfluence parameter was set to 1.0, giving full weight to information propagated from neighboring nodes during the iterative aggregation process.

Finally, **GraphSAGE** algorithm-based node embedding was generated using the Neo4j Graph Data Science library functions. The algorithm was applied to the subgraph projection using the previously trained model identified taking the node degrees as input features. The GraphSAGE model was trained on the projected graph’. Only the node degree feature property was used as the input feature, and embedding dimensions of (16, 32, 64 and 128) were specified. The ‘mean’ aggregator was applied to combine information from sampled neighbors, and the ‘ReLu’ activation function was applied in the neural network. Neighborhood sampling was configured with sampleSizes: [25, 10], whereby up to 25 neighbors were sampled at the first layer and 10 neighbors at the second layer. A batchSize of 100 was used, five training epochs were performed, and maxIterations: 10 was specified to limit the optimization iterations. A learningRate of 0.01 was applied to control the magnitude of parameter updates, while a negativeSampleWeight of 20 was assigned to negative samples during training. A searchDepth of 5 was specified for the search procedure, and no L2 regularization was applied because penaltyL2 was set to 0.0. A tolerance of 1e-4 was used as the convergence/termination criterion. In all methods above, a random seed of 42 was specified to ensure reproducibility of the stochastic feature generation.

### MOSAIC embedding generation

Given a graph adjacency matrix (*A*) of the underlying graph, MOSAIC first constructs a row-normalized transition operator:

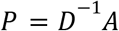

where (*D*) is the diagonal degree matrix.

The diffusion operator used in all reported experiments was the finite multi-step random-walk operator with α = 0. 9:

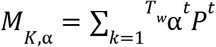

The experiments reported here used:

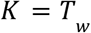

Therefore, the operator applied to a vector (x) was computed as:

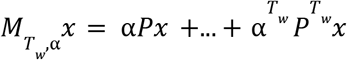

The diffusion operator was applied in matrix-free form using the Krylov-Ritz solver rather than explicitly constructing (*M*_*K*_).

The Krylov solver (**Algorithm 1** pseudocode) uses a single normalized random starting vector:

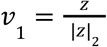

where (*z*) is a random vector sampled from a standard normal distribution. A single-vector Arnoldi procedure was used to construct the Krylov basis. At each iteration, the basis vectors were orthogonalized using modified Gram-Schmidt re-orthogonalization. The maximum Krylov dimension was set equal to the requested embedding dimension plus a constant *q*:

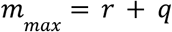

Thus, the implementation use an oversampled candidate space; at completion, the Arnoldi procedure provides at most (*r* + *q*) Ritz components, and all available final Ritz components are retained for the (*r* + *q*)-dimensional embedding. Nevertheless, the convergence scores described below determine the ordering of the retained embedding coordinates rather than performing an additional truncation from a larger candidate set, however oversampling provides a means to refine the ordered list even further.

The numerical parameters used in all experiments were:

1. Krylov dimension: (*m* _*max*_ = *r* + *q*, *q* = 1)
2. convergence tolerance: (10 *−*6)
3. single-vector Arnoldi initialization
4. normalized random starting vector

At every Arnoldi iteration, the projected Hessenberg matrix was formed and its eigendecomposition was computed to obtain the current Ritz values. The sequence of Ritz values across iterations was retained to monitor their numerical stability throughout the Krylov process. Following completion of the Arnoldi iterations, the final Hessenberg matrix was decomposed to obtain the final Ritz values and corresponding Ritz vectors.

Unlike conventional spectral truncation based solely on eigenvalue magnitude, MOSAIC orders the available spectral components according to their convergence stability during the Arnoldi iterations.

For each final Ritz value 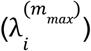 a convergence score was computed as the average distance between that final Ritz value and the closest Ritz value at each preceding Arnoldi iteration:

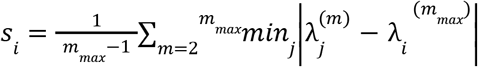

For complex-valued Ritz values, the modulus of the difference is used. Thus, the score measures the average distance between each final Ritz value and its nearest value at the preceding Arnoldi dimensions; it is not a measure of successive changes between adjacent iterations. Lower convergence scores indicate that a final Ritz value was more consistently approached during the Krylov process. The final Ritz components are ordered by increasing convergence score, with the corresponding Ritz vectors providing the node embedding coordinates. It should be noted that the Krylov-Ritz naturally tend to converge to the spectrally isolated eigenvalues with the fastest convergence rate.

Because (*m*_*max*_ = *r* + 1) in the present implementation, an oversampling parameter is used, and the number of retained Ritz components is the number of available final Ritz pairs plus 1 (up to (*r* + 1)). The implementation therefore applies a separate residual-based tolerance to discard Ritz pairs, and it performs an additional selection from more than (= *r*) candidates.

### Embedding construction

The selected Ritz vectors were transformed into node embeddings using:

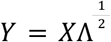

where (*X*) denotes the selected Ritz vectors and (Λ) contains the corresponding Ritz values.

Each reconstructed Ritz vector was subsequently normalized to unit Euclidean norm, ensuring a consistent scale across embedding dimensions. The resulting normalized Ritz vectors were used directly as the node embedding coordinates for downstream prediction tasks.

The current implementation evaluates MOSAIC as a diffusion-based graph representation method operating on an aggregated biomedical graph projection. Although the input data originate from a knowledge graph, relation types and ontology predicates are not explicitly modeled in the present formulation. Therefore, the reported experiments assess topology-based representation learning rather than relation-aware knowledge graph embedding (**Algorithm 2**).

### Node embedding classification

To evaluate the quality of the learned node representations, we performed a downstream node label prediction task using the generated embeddings as input features. The objective of this analysis was to assess whether the embedding coordinates preserve information relevant for distinguishing node classes without providing the classifier with additional graph structural information. UMAP projections of the 16 dimensional embeddings were computed using the R package uwot v0.2.3 [21].

For each embedding method, node representations were generated at four embedding dimensions (*r* = **16, 32**,**64**, and **128**). The resulting embedding matrix:

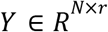

was used as the feature matrix, and the corresponding node labels were used as classification targets.

To avoid information leakage during model selection, the labeled nodes were randomly divided into three mutually exclusive subsets: training (**70%**), validation (**10%**), and testing (**20%**). The training subset was used to fit the classifier parameters, the validation subset was used for early stopping and model selection, and the test subset was used exclusively for final performance evaluation. Binary classification was performed using the “xgboost” package (v1.7.11.1 [22]) with a gbtree booster, logistic objective (binary:logistic), AUC evaluation metric, learning rate (eta) of **0.1**, maximum tree depth of **5**, and subsampling rates of **0.8** for both rows and columns; the model was trained for a maximum of **100** boosting rounds with early stopping after 10 rounds without improvement in validation AUROC.

Test set predictions were thresholded at 0.5 probability to obtain binary outputs. Model performance was evaluated using base R for accuracy and confusion matrix, and the “pROC” package (v1.19.0.1 [23]) for receiver operating characteristic (ROC) curve computation and area under the curve (AUC); results were visualized via plot.roc(). The analysis was repeated 10 times for each node embedding method and embedding dimension through random partitioning of the data in training, validation and test subsets and the mean and standard deviation of the derived AUCs were reported. Other auxiliary R packages were incorporated in the analysis including “Matrix” (v1.7-0) [24] and “RSpectra” (v0.16-1) [25].

The training subset was used exclusively for classifier fitting, while the test subset was reserved as an independent evaluation set. No model parameters, embedding configurations, or training decisions were selected using the test data. The reported AUROC values therefore represent performance on previously unseen samples after all methodological choices had been fixed. The use of a held-out test set was chosen to provide a direct assessment of the generalization capability of the learned representations while avoiding information leakage between training and evaluation stages. The classifier was trained using a fixed number of boosting iterations determined prior to evaluation. Because no adaptive model selection was performed using the test data, the held-out test set remained independent and was used only for final performance assessment.

### Robustness to Edge Removal Methods

To evaluate the robustness of MOSAIC to progressive loss of graph connectivity, we constructed a heterogeneous graph containing two entity types, denoted as Gene and Phenotype. Node-label prediction was performed using embeddings generated by MOSAIC and, for comparison, FastRP, a topology-based graph embedding method. The evaluation was designed to quantify the extent to which each embedding method preserves information relevant to node-label prediction as the observed graph becomes increasingly sparse. We generated a sequence of perturbed graphs by randomly removing increasing fractions of the original edges. Specifically, the proportion of removed edges was varied from 0 to 0.99 in increments of 0.09, with the removed-edge fraction. For each perturbed graph, embeddings were independently learned from the remaining graph structure using MOSAIC and FastRP. These embeddings were subsequently used as input to the same node-label prediction procedure, thereby ensuring that differences in predictive performance were attributable primarily to the information retained by the respective embedding representations rather than to differences in the downstream prediction model. Robustness was assessed using the mean area under the receiver operating characteristic curve (AUROC). Mean AUROC measures the probability that a randomly selected positive instance receives a higher prediction score than a randomly selected negative instance and is therefore particularly suitable for evaluating ranking-based prediction performance independently of a particular classification threshold.

For each edge-removal level, mean AUROC was calculated using the node-label predictions generated from the corresponding embeddings after 3 edge removal repeats. The same edge-removed graphs were used for MOSAIC and FastRP embedding. Higher AUROC values indicate greater preservation of label-discriminative information under the specified level of graph perturbation. To further characterize the dependence of embedding robustness on edge type, the perturbation analysis was conducted under four distinct edge-removal schemes: (A) removal of edges of all types, (B) removal restricted to Gene-Phenotype edges, (C) removal restricted to Gene-Gene edges, and (D) removal restricted to Phenotype-Phenotype edges. Thus, the first analysis measures overall robustness to graph sparsification, whereas the latter three analyses isolate the contribution of inter-type and intra-type connectivity. For each perturbation scheme, MOSAIC and FastRP were evaluated on identical edge-removed graphs and using the same node-label prediction protocol, enabling a direct comparison of their sensitivity to structural information loss.

### Local deviation analysis methods

To quantify local geometric consistency, we computed a node-level local deviation score for each embedding method and embedding dimensionality. For each node *v*, we first identified its immediate neighborhood *N*(*v*) according to the graph connectivity used throughout the analysis and computed the centroid of the corresponding neighbor embeddings. The local deviation of *v* was then defined as the distance between its embedding vector and this neighborhood centroid, where *z*_*v*_ denotes the embedding of node *v* and ∥ · ∥ _2_ is the Euclidean norm. Thus, smaller values indicate greater alignment between a node and the embeddings of its immediate neighbors, whereas larger values indicate greater local geometric deviation. We calculated this measure independently for each of the five embedding methods and for embedding dimensions of 16, 32, 64, and 128. The resulting node-level deviation scores were summarized as empirical distributions for each method-dimension combination and visualized in Figure 5. To facilitate comparison across methods and dimensionalities, the same definition of neighborhood, distance metric, and distributional representation was applied throughout. We examined the resulting distributions descriptively in terms of their central tendency, concentration near zero, dispersion, skewness, tail behavior, and, where applicable, multimodality. These analyses were intended to characterize differences in the local geometry induced by the embedding methods rather than to provide an intrinsic measure of embedding quality.

### Predicting links between genes and phenotypes

To evaluate whether MOSAIC node embeddings preserve biologically meaningful relationships between nodes, we performed a gene-phenotype link prediction experiment. An induced biomedical graph projection containing **5**,**171** human genes and **10**,**147** phenotypes was used as the evaluation network. Because the objective of this experiment was to evaluate topology-preserving graph representations, the available biomedical relationships were aggregated into a graph projection prior to embedding generation. The resulting representation captures structural connectivity patterns but does not explicitly encode relation-specific semantics. To prevent information leakage, observed gene-phenotype associations were partitioned before embedding generation: training (**70%**), validation (**10%**), and testing (**20%**). The graph used for computing node embeddings contained only the training edges. Validation and test edges were removed from the graph prior to applying MOSAIC and were used exclusively for downstream evaluation. For negative examples, disconnected gene-phenotype pairs were sampled from the graph and screened for the absence of a direct link. A balanced dataset comprising ∼**1**,**100**,**000** gene-phenotype node pairs was created for the link prediction task. Half of the pairs corresponded to existing gene-phenotype associations present in the induced subgraph (positive examples), while the remaining half represented node pairs without a direct connection (negative examples were ∼400,000 edges of genes-phenotypes made of genes not expressed in the tissues with the indexed phenotype, and double checked w.r.t. the DDKG graph).

For each node pair (*u, v*) the corresponding node embedding vectors were concatenated:

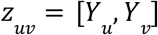

The resulting pairwise representations were used as input features for an XGBoost binary classifier. Classifier training used only training pairs, while validation pairs were used for early stopping. Final link prediction performance was evaluated on the independent test pairs using AUROC.

### DICER1 pleiotropy

To assess the performance and biological relevance of the graph-based link-prediction framework, we performed a focused analysis of DICER1, a well-characterized pleiotropic disease-associated gene. The trained XGBoost classifier was applied to the DICER1 embedding and each of the 10,147 phenotype nodes in the graph to generate an estimated edge probability for every DICER1-phenotype pair. Predicted probabilities were compared with the 137 phenotypes having documented direct DICER1 edges in the underlying graph, with P > 0.90 used as a descriptive high-confidence threshold. We also examined the complete distribution of predicted probabilities across all phenotypes and quantified predictions exceeding P > 0.99 . The biological relevance of the resulting predictions was assessed in the context of established DICER1 genotype-phenotype associations and its recognized broad phenotypic spectrum. Because the graph embeddings were derived from the knowledge graph containing known associations, recovery of existing DICER1-phenotype edges was interpreted as evidence of internal concordance, rather than independent validation; predictions for phenotypes without established edges were considered hypothesis-generating and requiring external validation.

## Results

### Graph topological characteristics

The human gene-phenotype graph, comprising 5,171 human genes and 10,147 human phenotypes (15,318 nodes in total), exhibits a globally compact, locally clustered, and strongly community-structured topology. Its mean shortest path of 2.9486 and diameter of only 6 indicate highly efficient global connectivity, with most nodes separated by roughly three steps and even the most distant nodes requiring no more than six steps. The mean eccentricity of 5.021±0.519, relatively small compared with the diameter, further indicates that most nodes occupy similarly compact positions within the network and that there are few extremely peripheral or elongated regions. The transitivity of 0.280±0.295 demonstrates substantial local interconnectedness, while the standard deviation being approximately as large as the mean reveals pronounced heterogeneity in clustering, suggesting that some regions form densely interconnected biological neighborhoods whereas others are comparatively sparse and may contain peripheral or bridging nodes. The presence of two exceptionally large 122-node cliques provides evidence for highly dense network cores, although their biological interpretation depends on the graph construction. The harmonic centrality of 5500.185±744.111 indicates meaningful variation in global accessibility, with a coefficient of variation of approximately 13.5%, suggesting that the network contains a broad population of relatively central nodes rather than being controlled exclusively by a tiny set of hubs; such nodes may correspond to pleiotropic genes, broadly associated phenotypes, or connector entities, although network centrality should not be equated directly with biological importance because annotation and study biases can strongly affect these measures. The identification of 14 Louvain communities further demonstrates clear mesoscale organization, potentially corresponding to distinct functional, disease, or phenotypic modules, although the number of communities alone is insufficient and should be accompanied by the actual modularity score, community sizes, gene/phenotype composition, internal density, pathway enrichment, and inter-community connectivity. Taken together, the combination of short path lengths, small diameter, relatively uniform eccentricity, substantial but heterogeneous transitivity, exceptionally dense cores, and 14 communities is consistent with a hierarchical and potentially small-world-like biological architecture in which specialized, highly clustered gene-phenotype modules are embedded within an efficiently interconnected global network. Overall, the current statistics strongly support the interpretation of the network as a globally efficient yet locally heterogeneous and modular human gene-phenotype system (**Table 1**).

**Table 1.** Gene-phenotype graph cumulative topological metrics.

| Graph Metric | Value |
| --- | --- |
| Mean Shortest Path | 2.948559 |
| Diameter | 6 |
| Eccentricity | 5.020956±0.5189038 |
| Transitivity | 0.2803142±0.2947593 |
| Largest Cliques | 122 Nodes, Count: 2 |
| Harmonic Centrality | 5500.185±744.1109 |
| Louvain Clusters | 14 |

### Theoretical computational complexity

The proposed MOSAIC method is spectral and deterministic, relying primarily on repeated sparse matrix-vector multiplications and an iterative eigendecomposition, making it comparable in efficiency to spectral methods and significantly cheaper than deep neural or walk-based methods. Unlike Node2Vec or GraphSAGE, it avoids stochastic optimization and deep model training, while being more expressive than FastRP due to the multistep diffusion operator (**Table 2**). The memory requirements of graph node embedding methods depend strongly on whether the method performs explicit spectral computation, random-walk sampling, or neural message passing. MOSAIC requires storage of the sparse graph, the Krylov basis, and the reduced spectral representation, resulting in (*O*(|*E*| + *nr* + *r*^2^)) memory complexity. FastRP has a comparable asymptotic requirement of (*O*(|*E*| + *nr*)) because it stores only sparse graph information and projected node representations. Node2Vec additionally requires storage for sampled random walks, giving (*O*(|*E*| + *nwL* + *nr*)) complexity, where (*w*) and (*L*) denote the number and length of walks. GraphSAGE introduces additional training memory requirements because intermediate activations and sampled computation graphs must be retained for back-propagation, resulting in approximately (*O*(|*E*| + *nd* + *BS*^*L*^*r*)) memory during training. HashGNN avoids full neural message passing and has memory requirements closer to conventional embedding approaches, approximately (*O*(|*E*| + *Lnr*)). Therefore, MOSAIC maintains sparse graph scalability while avoiding the potentially large sampling or activation-storage overheads associated with Node2Vec and GraphSAGE.

**Table 2.** Comparison of the theoretical computational complexities of MOSAIC and major graph embedding methods ***n* =** |***V***|: number of nodes, |***E***|: number of edges, ***r***: embedding dimension, ***N***: number of GNN layers, ***l***: random walk length, ***w***: number of walks per node, ***n***_***w***_: Skip-Gram context window size, ***n***_***h***_ : hashed/sampled neighbors per node.

| Method | Core Mechanism | Time Complexity (approx.) | Memory Complexity | Scalability & Remarks |
| --- | --- | --- | --- | --- |
| **MOSAIC** | Eigen-decomposition of multistep random-walk operator | $O(\vert E\vert \cdot r + n \cdot r^2)$ for operator applications and top-$r$ eigenvectors (using iterative eigensolver) | $O(\vert E\vert + n \cdot r + n \cdot r^2)$ | Efficient for sparse graphs; scales linearly in edges per diffusion step; no stochastic training required. |
| **FastRP [26]** | Random projections of normalized powers of adjacency | $O(\vert E\vert \cdot r)$ | $O(\vert E\vert + n \cdot r)$ | Extremely fast; suitable for billion-scale graphs; closed-form linear operator. |
| **Node2Vec [1]** | Biased random walks + SkipGram training | $O(l \cdot w \cdot n_w + m \cdot r) \approx O(\vert E\vert \cdot w)$ | $O(\vert E\vert + n \cdot r + l \cdot w \cdot n_w)$ | High cost due to random-walk sampling and SGD training; not ideal for very large graphs. |
| **GraphSAGE [27]** | Aggregation of neighbor features via neural layers | $O(N \cdot \vert E\vert \cdot r)$ (for $N$ layers) | $O(\vert E\vert + n \cdot r)$ | Scales linearly with graph size; higher cost with more layers and feature dimensions. |
| **HashGNN [28]** | Hash-based neighbor sampling and aggregation | $O(N \cdot n_h \cdot r)$, where $n_h \ll \vert E\vert $ | $O(\vert E\vert + n \cdot r)$ | Reduces aggregation cost by compressed hashing; faster variant of GraphSAGE. |

### Node label prediction

**Figure 1.** shows node type classification AUROC across damping factors of 0.1, 0.5, and 0.9, embedding dimensions from 2 to 128, and diffusion lengths from 2 to 32, with three repeated analyses per setting. AUROC increased with embedding dimension and approached 1.0 across the tested parameter combinations. In the 32 dimensional comparison shown in **Figure 2**, MOSAIC achieved an AUROC of 0.9902, compared with 0.8766 for the largest magnitude spectral embedding. UMAP projections of the 16 dimensional embeddings showed clearer separation of gene and phenotype nodes with MOSAIC than with FastRP **(Figure 3)**.

**Figure 1.**
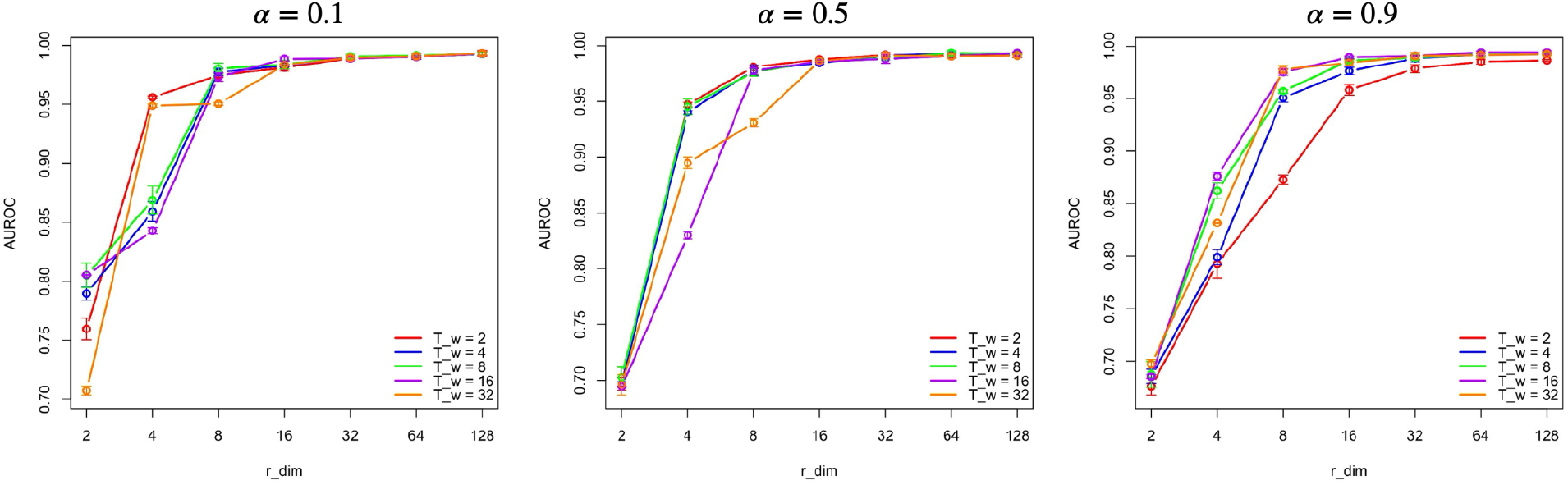
The mean AUROC of the node label (gene or phenotype) prediction as a function of MOSAIC node embedding dimensions, damping factor, and diffusion length for the projected gene-phenotype graph

**Figure 2.**
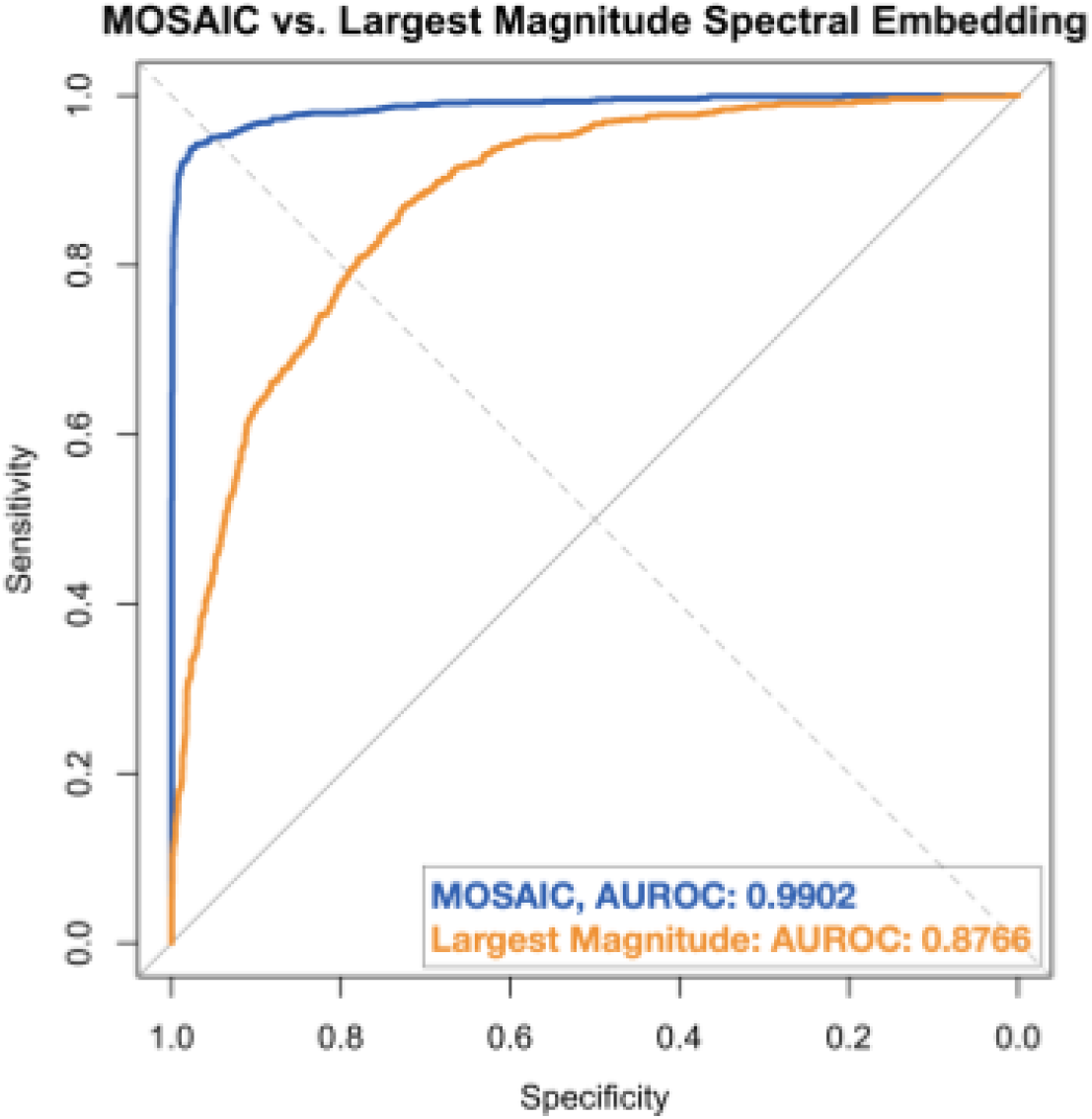
Comparison of the MOSAIC-and largest magnitude eigendecompostion-based node label predictions of example one using AUROCs.

**Figure 3.**
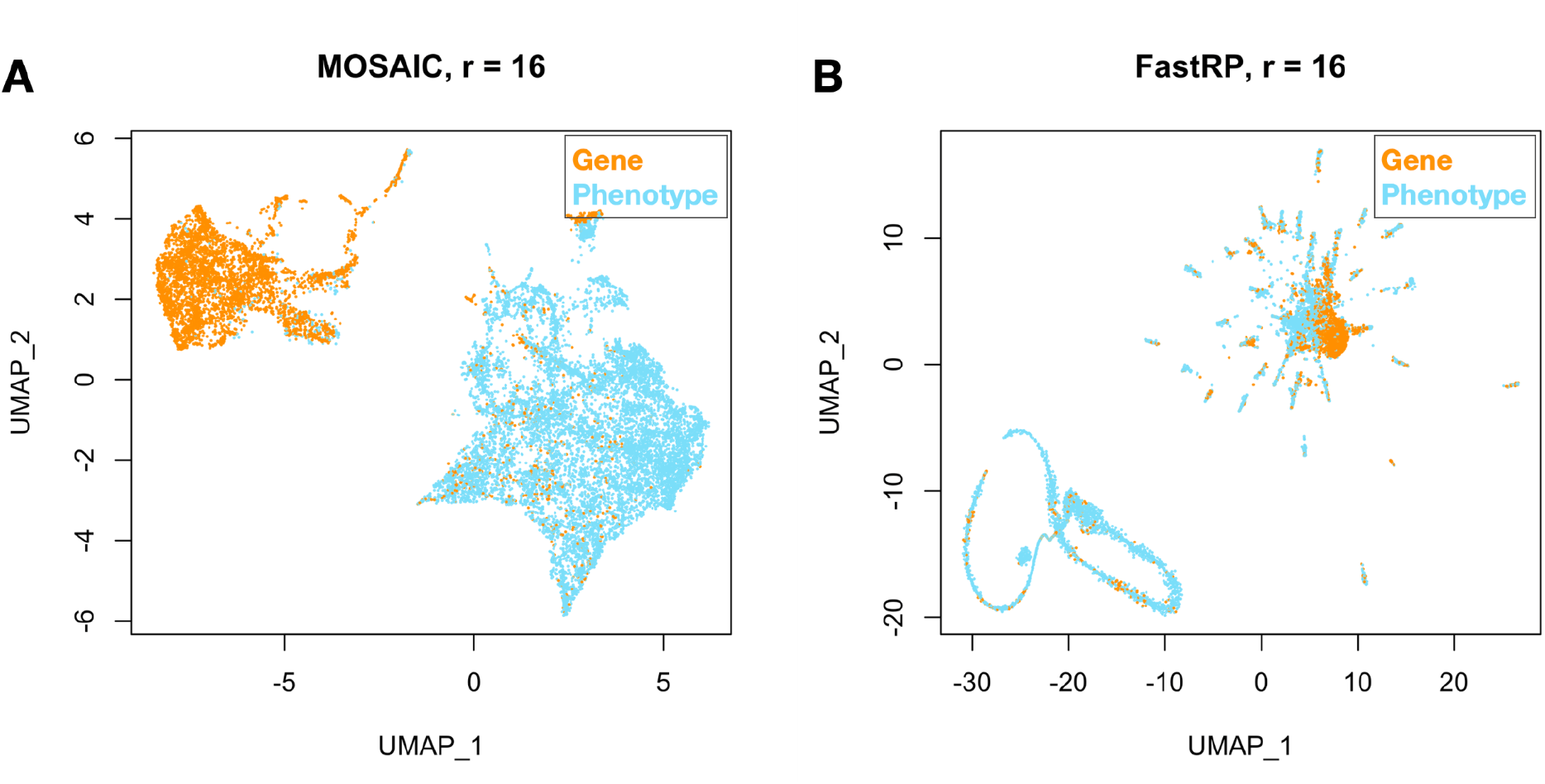
UMAP representation of the human genes and phenotypes embeddings derived from the **A)** MOSAIC transformation and **B)** FastRP of the example subgraph with r = 16.

**Table 3** compares node type classification performance across embedding dimensions of 16, 32, 64, and 128. Across 10 repeated training, validation, and test partitions, MOSAIC achieved mean AUROCs of 0.9863 ± 0.0019, 0.9903 ± 0.0013, 0.9916 ± 0.0010, and 0.9926 ± 0.0013, respectively. MOSAIC exceeded all comparison methods at every tested dimension. Node2Vec was the strongest comparator, with mean AUROCs ranging from 0.9547 to 0.9768, while FastRP increased from 0.9217 to 0.9710 as embedding dimension increased. HashGNN and GraphSAGE showed lower performance across all dimensions. These results indicate that MOSAIC embeddings retain information sufficient to distinguish gene and phenotype nodes across the evaluated embedding dimensions.

**Table 3.** Comparing node type classification AUC as a function of embedding dimensions.

| Method | r = 16 | r = 32 | r = 64 | r = 128 |
| --- | --- | --- | --- | --- |
| **MOSAIC** | **0.9863±0.0019** | **0.9903±0.0013** | **0.9916±0.001** | **0.9926±0.0013** |
| FastRP | 0.9217±0.003 | 0.9539±0.0031 | 0.9615±0.0044 | 0.971±0.0031 |
| Node2Vec | 0.9547±0.0032 | 0.9725±0.0032 | 0.9763±0.003 | 0.9768±0.003 |
| HashGNN | 0.7977±0.0088 | 0.8044±0.0058 | 0.8091±0.0054 | 0.8064±0.0067 |
| GraphSAGE | 0.8265±0.005 | 0.8337±0.0075 | 0.8346±0.0061 | 0.8364±0.0082 |

### Robustness to edge removal

To evaluate the robustness of MOSAIC to progressive loss of graph structure, we systematically removed increasing proportions of existing edges from the graph and evaluated node-label prediction using embeddings learned from each perturbed graph. Specifically, edge-removal ratios ranging from 0 to 0.99 in increments of 0.09 were considered, and the resulting node-label prediction performance was quantified using the area under the receiver operating characteristic curve (AUROC). FastRP was used as a reference embedding method. The analysis was performed under four perturbation regimes: removal of all edge types, removal of only Gene-Phenotype edges, removal of Gene-Gene edges, and removal of Phenotype-Phenotype edges. This experimental design allows robustness to be assessed not only with respect to overall graph sparsification, but also with respect to the selective loss of inter-type and intra-type relational information. Removal of all edges.

As shown in **Figure 4A**, MOSAIC consistently achieves higher AUROC than FastRP across the entire range of edge-removal ratios. Both methods exhibit a gradual reduction in node-label prediction performance as the graph becomes increasingly sparse, which is expected because edge removal progressively reduces the relational information available to the embedding algorithms. However, the degradation is substantially less pronounced for MOSAIC, resulting in an increasingly larger performance gap relative to FastRP as the fraction of removed edges increases. Notably, when 99% of the original edges have been removed, MOSAIC retains an AUROC of approximately 0.75, whereas FastRP decreases to below 0.60. This indicates that the representations generated by MOSAIC preserve information relevant to node-label discrimination even under extreme structural sparsification. From a representation-learning perspective, this behavior suggests that MOSAIC is able to retain more stable node-level signals when local neighborhood information becomes progressively unavailable, potentially through a more effective integration of higher-order structural information and/or more robust aggregation of the remaining relational context. Removal of Gene-Phenotype edges.

**Figure 4.**
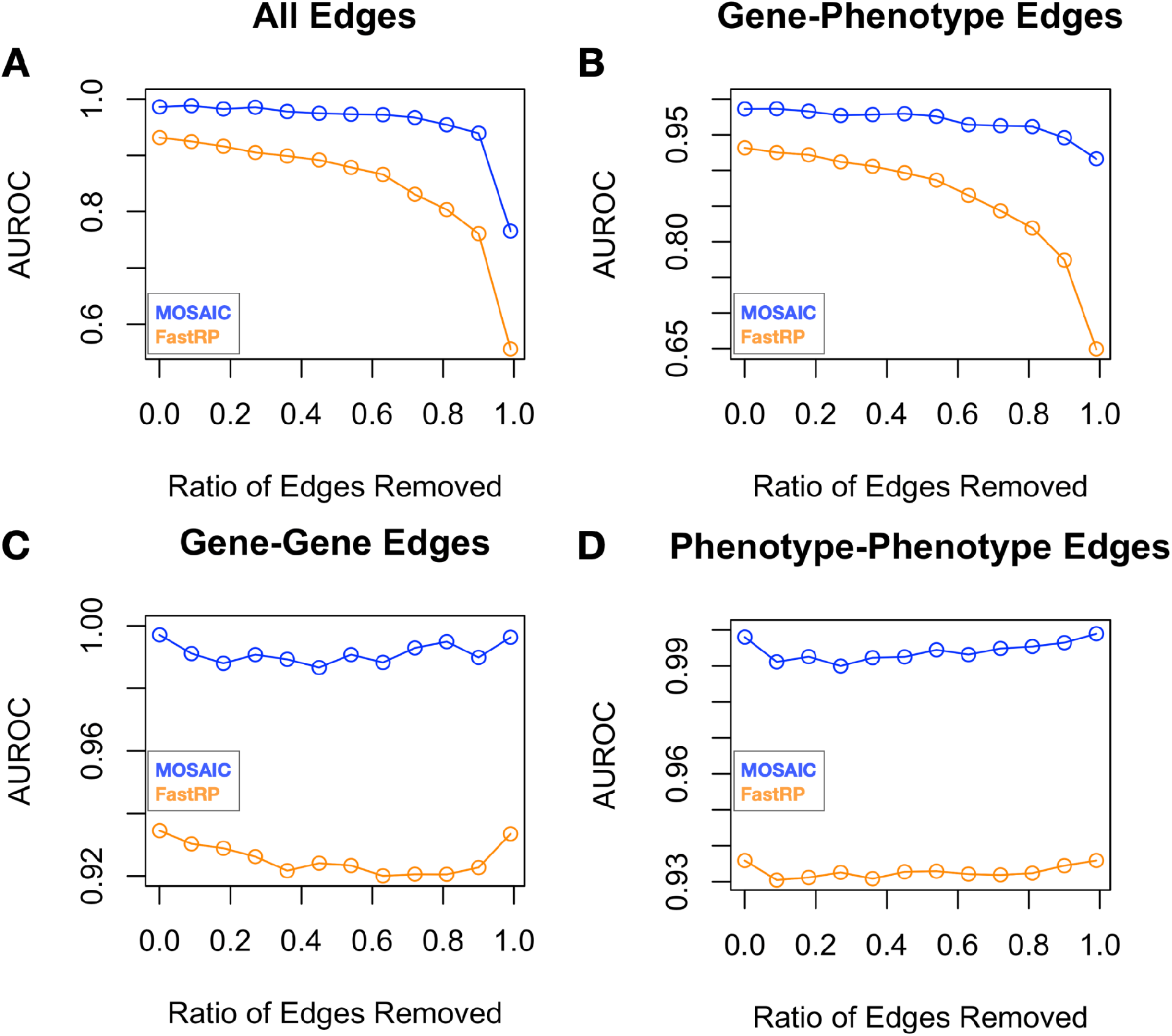
Robustness of MOSAIC and FastRP compared together (r = 16) as measured by AUROC of label prediction task with edge removal ratio ranging 0 to 0.99 from **A)** all edges, **B)** Gene-Phenotype edges, **C)** Gene-Gene edges and **D)** Phenotype-phenotype edges.

**Figure 5.**
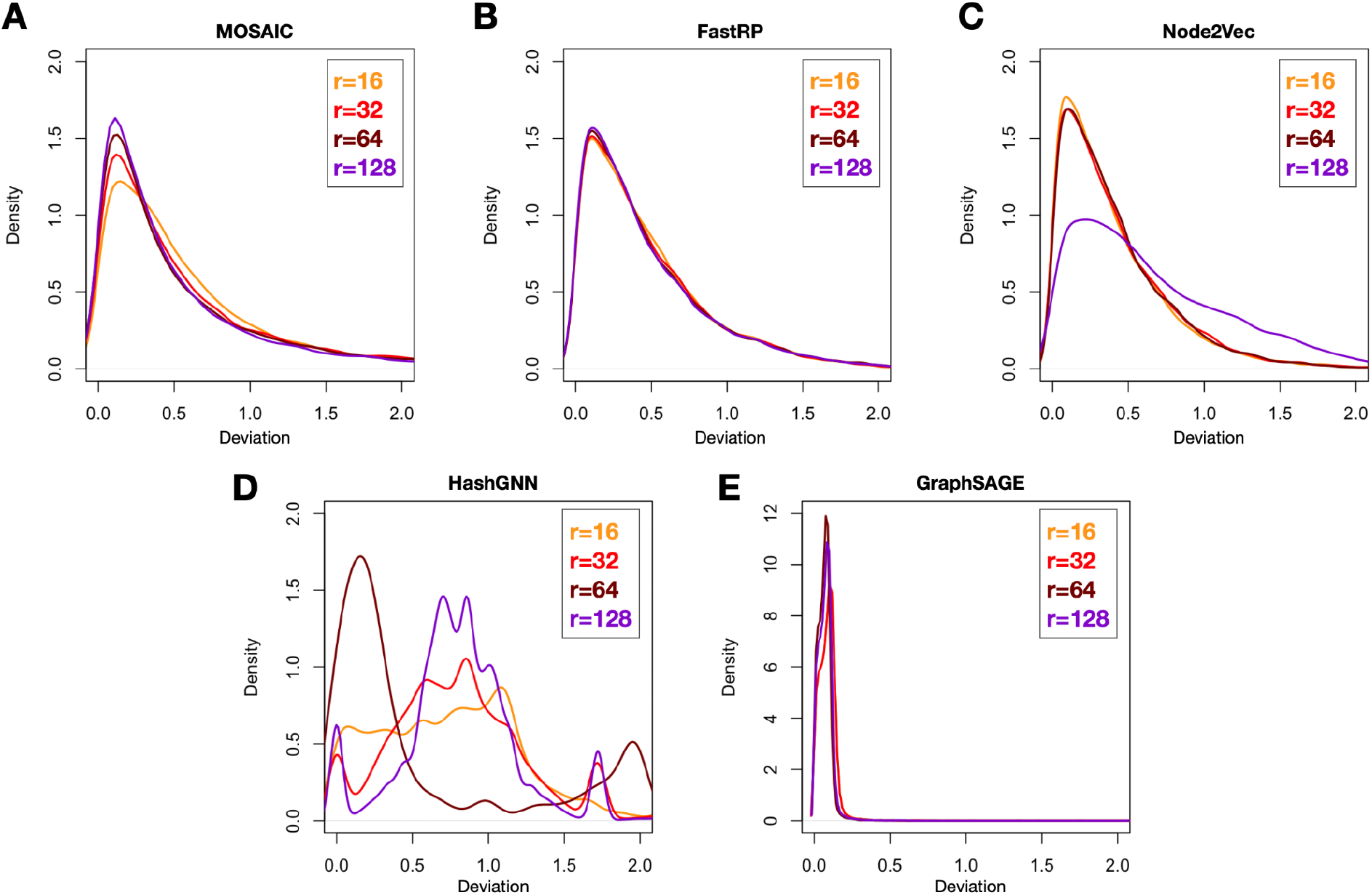
Local deviation analysis comparing MOSAIC with FastRP, Node2Vec, HashGNN and GraphSAGE. The deviation reveals how smoothly the embeddings transition from node to node traversing across the graph.

**Figure 4B** shows a similar, but more pronounced, pattern when the perturbation is restricted to Gene-Phenotype edges. MOSAIC maintains higher AUROC than FastRP at every edge-removal ratio, with the difference between the two methods increasing as the proportion of removed Gene-Phenotype edges grows. The performance of both methods decreases with increasing edge removal, confirming that the inter-type connections provide important information for the node-label prediction task. Nevertheless, MOSAIC remains substantially more resilient to the loss of these connections. At 99% removal of Gene-Phenotype edges, MOSAIC achieves an AUROC of approximately 0.85, compared with an AUROC below 0.65 for FastRP. The stronger absolute performance of MOSAIC under this perturbation indicates that its embedding space retains discriminative information even when the principal connections between the two entity types are almost completely eliminated. This robustness is particularly relevant for heterogeneous graphs, where removal of inter-type edges can cause a rapid loss of cross-type context and consequently fragment the information flow between the two node populations. The results suggest that MOSAIC can compensate for this loss more effectively than FastRP, potentially because its representations encode structural dependencies that are less exclusively dependent on direct Gene-Phenotype connectivity. Removal of Gene-Gene edges.

The results in **Figure 4C** further demonstrate the robustness of MOSAIC when only Gene-Gene edges are removed. MOSAIC consistently outperforms FastRP throughout the perturbation range, with the performance gap increasing as the graph is progressively deprived of Gene-Gene connections. Both methods initially exhibit a decline in AUROC as the fraction of removed Gene-Gene edges increases, reaching a minimum at approximately 36% edge removal. Interestingly, beyond this point, the prediction performance increases despite further removal of Gene-Gene edges. At 99% removal, MOSAIC achieves an AUROC of approximately 0.98, while FastRP remains around 0.93. The non-monotonic response indicates that the relationship between graph sparsity and predictive performance is not necessarily linear, particularly when a specific edge type is selectively perturbed. One possible explanation is that removing a large fraction of Gene-Gene edges changes the relative contribution of the remaining edge types, effectively increasing the structural or statistical prominence of the remaining Gene-Phenotype and/or Phenotype-Phenotype connections. If these remaining relations provide stronger or less noisy information for the target labels, their increased relative contribution can partially or fully compensate for the loss of Gene-Gene connectivity. The particularly high AUROC observed at extreme Gene-Gene removal also suggests that Gene-Gene edges are not indispensable for the underlying node-label discrimination task, and that MOSAIC can exploit alternative structural signals when this relationship type is largely absent. Removal of Phenotype-Phenotype edges.

A comparable pattern is observed for selective removal of Phenotype-Phenotype edges in **Figure 4D**. MOSAIC consistently exhibits higher AUROC than FastRP, and the performance difference becomes increasingly pronounced as the proportion of removed Phenotype-Phenotype edges increases. Both methods initially show decreasing prediction performance, with the decline continuing until approximately 27% Phenotype-Phenotype edge removal. Beyond this point, however, AUROC increases despite further removal of Phenotype-Phenotype edges. At 99% Phenotype-Phenotype edge removal, MOSAIC reaches an AUROC of approximately 0.99, compared with approximately 0.94 for FastRP. As in the Gene-Gene perturbation experiment, this non-monotonic behavior suggests that selective removal of a particular intra-type relationship can alter the balance of information encoded by the remaining graph structure rather than simply reducing the total amount of useful information. In particular, the remaining Gene-Phenotype and Gene-Gene relations may become comparatively more informative for node-label prediction once Phenotype-Phenotype connectivity is substantially reduced. The near-perfect AUROC obtained by MOSAIC under this extreme perturbation further indicates that the information required for the prediction task can be recovered from structural signals other than Phenotype-Phenotype connectivity. Overall, the four perturbation experiments demonstrate that MOSAIC maintains substantially more stable predictive performance than FastRP under progressive graph sparsification. The advantage is observed for both global edge removal and targeted removal of individual relationship types, and the performance gap generally becomes larger as more structural information is removed. Importantly, the results also show that predictive performance cannot be interpreted solely as a function of the total number of remaining edges. In the Gene-Gene and Phenotype-Phenotype perturbation experiments, extreme removal of a specific relationship type is followed by a recovery of AUROC, indicating that the composition of the remaining graph structure can be more important than its overall density. Taken together, these findings support the interpretation that MOSAIC produces embeddings that are less dependent on any single edge type and are therefore more robust to structural perturbations. The substantially higher AUROC retained under severe edge removal suggests that MOSAIC captures transferable or higher-order structural information that remains accessible even when direct neighborhood information is heavily degraded.

### Local deviation analysis

**Figure 5.** provides detailed information on the distribution of local deviations across five embedding methods and four embedding dimensions. The local deviation distributions can be interpreted as characterizing the local geometric consistency induced by each embedding method, where the distance for a node measures the deviation of its embedding from the centroid of its immediate neighborhood. Across the five methods, MOSAIC and FastRP exhibit broadly similar behavior, with strongly right-skewed distributions, a high concentration of nodes at small node-to-neighborhood distances and a comparatively sparse but pronounced population of nodes with large deviations. This combination of a mean close to zero and a heavy right tail suggests that the embeddings preserve strong local proximity for most nodes while permitting a relatively small subset of structurally heterogeneous, boundary or weakly represented nodes to occupy substantially different locations from their neighbors. The progressive narrowing of MOSAIC as dimensionality increases from 16 to 128 indicates that additional dimensions appear to stabilize the local geometry and reduce heterogeneity in node-to-neighborhood distances, potentially reflecting a progressively more faithful representation of local structure. In contrast, the relative invariance of FastRP’s distributional diffuseness across dimensions suggests that its local geometric organization is comparatively insensitive to embedding dimensionality over this range; increasing representational capacity does not substantially alter the dispersion of neighborhood-consistency scores.

Node2vec presents a different dimensionality response: although its distributions remain right-skewed, increasing the embedding dimension produces progressively more diffuse distributions with a heavier right tail. This indicates that additional dimensions do not simply refine a common local geometry but instead permit greater differentiation among nodes, including an increasing population of nodes whose embeddings deviate substantially from the centroid of their neighborhoods. In this sense, Node2vec appears to use increased dimensionality to express greater heterogeneity in local structural roles, whereas MOSAIC uses increased dimensionality to produce greater geometric concentration.

HashGNN is qualitatively distinct from all three, exhibiting multimodal and irregular distributions with multiple local maxima whose locations and relative prominence change across dimensionalities. Such behavior suggests a more heterogeneous embedding geometry in which several distinct regimes of node-to-neighborhood consistency coexist; the lack of a stable unimodal form makes its response to dimensionality substantially less predictable and may indicate that different structural or semantic node populations are being separated differently as representational capacity changes.

Finally, GraphSAGE exhibits the strongest concentration near zero and essentially no substantial right tail, indicating that its node embeddings are highly aligned with the centroids of their immediate neighborhoods across the graph. The fact that this concentration changes little with dimensionality suggests that the local smoothness induced by the method is relatively dimension-invariant. Importantly, this should not automatically be interpreted as superior embedding quality: an extremely concentrated distribution can reflect effective preservation of local homophily, but it can also indicate over-smoothing or reduced local discriminability, particularly if embeddings of structurally distinct nodes become excessively similar.

Taken together, the five methods therefore appear to occupy different points on a spectrum between local geometric smoothness and local structural heterogeneity: GraphSAGE is the most strongly concentrated and locally smooth, MOSAIC becomes increasingly concentrated as dimensionality grows, FastRP maintains relatively stable local geometry across dimensions, Node2vec increasingly exposes heterogeneous local relationships at higher dimensions and HashGNN produces the most complex and dimension-dependent geometric structure. These observations are descriptive of the embedding geometry rather than evidence that one method is intrinsically better; determining which behavior is desirable requires relating these distributions to downstream objectives such as link prediction, node classification, community recovery preservation of structural roles.

### Predicting links between genes and phenotypes

To evaluate the impact of embedding dimensionality on the performance of graph-based link prediction models, we constructed an induced subgraph. This subgraph served as the benchmark dataset for assessing the ability of different graph embedding methods to capture biologically meaningful relationships. Node embeddings were generated using five graph representation learning approaches: MOSAIC, FastRP, Node2Vec, HashGNN, and GraphSAGE. For each method, embeddings were computed at four different dimensionalities (16, 32, 64, and 128 dimensions) to investigate how the size of the latent representation influences downstream predictive performance.

For each node pair, the embedding vectors of the source and target nodes were concatenated to generate the feature representation used for classification. These concatenated embeddings served as input to an XGBoost binary classifier trained to distinguish between connected and unconnected gene-phenotype pairs. After training, the classifier was applied to the independent test set to predict the presence or absence of direct associations. The predictive performance of each embedding method and embedding dimensionality was evaluated using the area under the receiver operating characteristic curve (AUROC). This analysis enabled a systematic comparison of the different graph embedding strategies and assessed the extent to which increasing embedding dimensionality improved the ability of the learned representations to recover known gene-phenotype associations. The resulting AUROC values provide a quantitative measure of the discriminative power of each embedding approach and identify the embedding dimensionalities that achieve the best link prediction performance (**Figure 6**).

**Figure 6:**
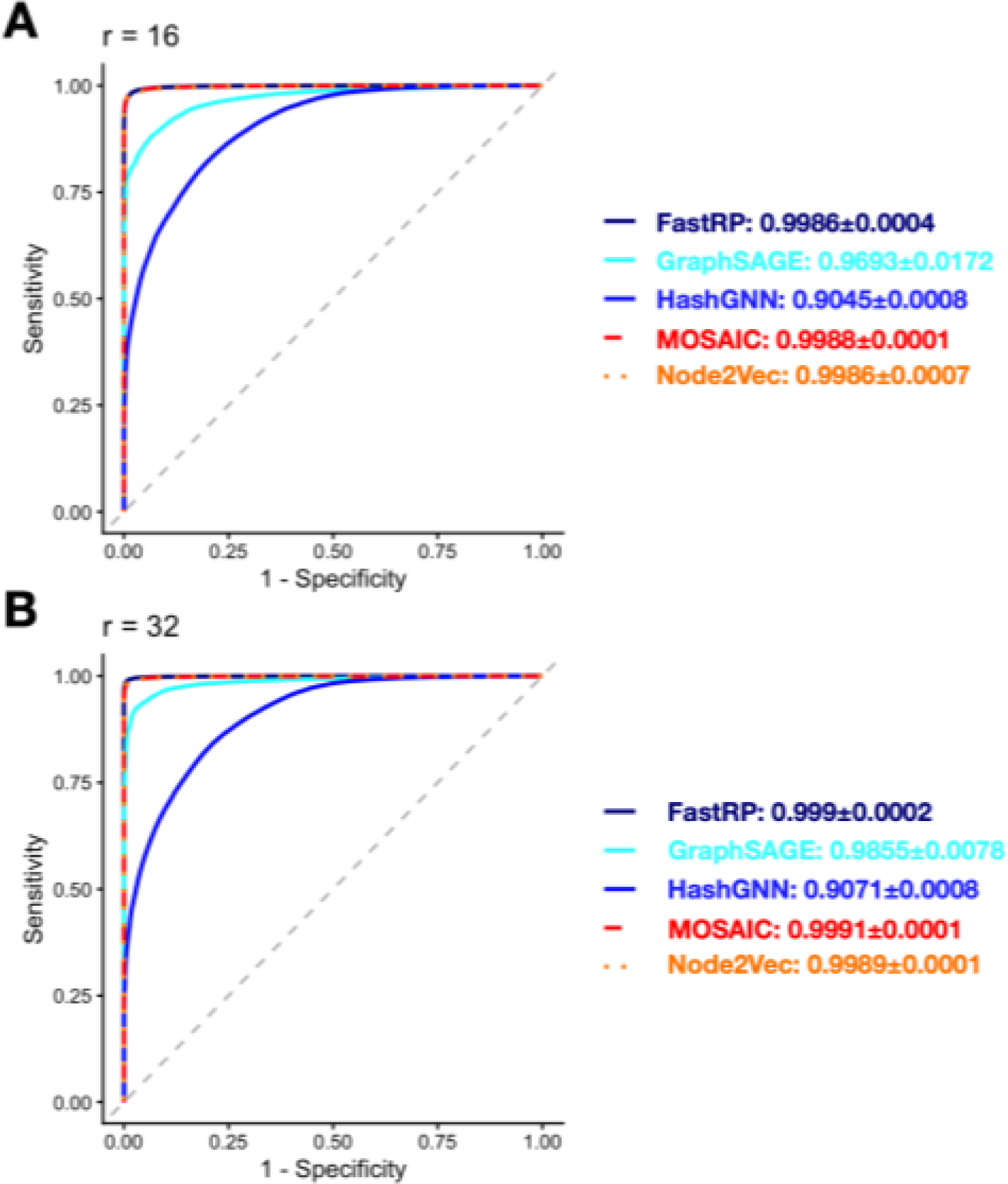
Comparison of AUAOC performance for MOSAIC and four baseline knowiliedge graph embedding methods (Node2Vec; FastRP, HashGNN, and GraphSAGE) ac1ross embedding dimensions (r=16 and 32). MOSAIC consis1enty achieves the highest AU AOC a1 every embedding dimension, with performance improving as embedding size increases,. demonstrating superior predictive accuracy robustness, and scalability,compared with the baseline methods (3 sampling to simulate repeats or each simulation)

### Link prediction use case: DICER1 pleiotropy

The model assigned high predicted probabilities to the majority of established DICER1-phenotype associations: 106 of 137 (77.4%) directly connected phenotypes had P > 0.90 (**Figure 7A**), indicating substantial concordance between the learned graph representation and known DICER1 biology. Across all 10,147 phenotypes, the predicted probability distribution was strongly shifted toward high values, with a prominent peak at approximately 0.98, and more than 600 phenotypes receiving P > 0.99 (**Figure 7B**). This pattern is consistent with the extensive phenotypic pleiotropy of DICER1, which is associated with manifestations spanning multiple organ systems and tumor types, and with previous work demonstrating that graph embeddings can capture both direct and indirect gene-phenotype relationships. The recovery of 77.4% of known associations supports the ability of the model to recognize established graph relationships, while the large number of very-high-probability phenotypes suggests that the model identifies a broad network of potentially relevant DICER1-phenotype relationships beyond those explicitly represented in the graph. However, the concentration of probabilities near 1 should not be interpreted as calibrated biological probabilities, particularly given the possibility of class imbalance and information leakage from graph-derived embeddings. Accordingly, the >0.99 predictions are best regarded as high-priority hypotheses for independent clinical, database, or experimental validation, rather than as confirmed novel associations.

**Figure 7:**
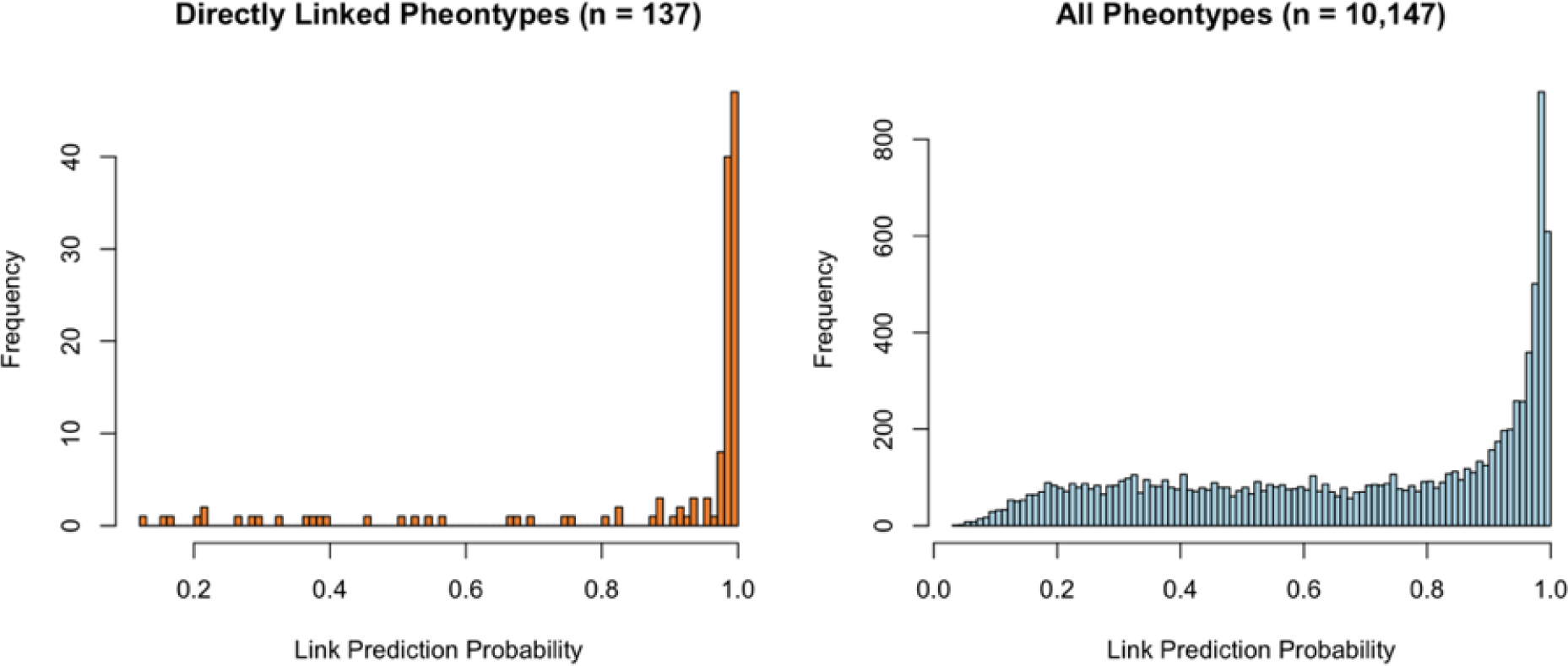
**A)** Distribution of link prediction scores between the human DICER1 gene and its directly linked phenotypes in the gene-phenotype graph. **B)** Predicted link probabilities of DICER1 and 10,147 human phenotypes.

The high recovery of known DICER1 associations is concordant with the established clinical literature. DICER1 syndrome is recognized as a pleiotropic tumor-predisposition disorder, with reported manifestations including pleuropulmonary blastoma, cystic lung lesions, cystic nephroma, rhabdomyosarcoma, thyroid follicular nodular disease, and Sertoli-Leydig cell tumors. The broad spectrum of phenotypes represented in current human phenotype resources further supports the observation that DICER1 has a relatively extensive phenotype neighborhood within a biomedical knowledge graph [29].

## Discussion

Graph embedding methods traditionally construct low-dimensional representations from eigenspaces associated with extremal eigenvalues of a graph-derived operator, such as the adjacency matrix, graph Laplacian, or normalized transition matrix. This ordering is motivated by asymptotic spectral theory, where eigenvalues of largest magnitude (or smallest nonzero Laplacian eigenvalues) characterize the dominant long-term behavior of the underlying dynamical system. While this approach provides strong theoretical guarantees, it implicitly assumes that spectral dominance alone determines the most informative subspace for representing graph structure. In practice, however, the contribution of an eigenmode to the action of an operator depends not only on its eigenvalue but also on its interaction with the subspace explored by the operator acting on data vectors.

The proposed framework instead emphasizes spectrally isolated eigenpairs that emerge as dominant under Krylov subspace iterations. From the perspective of numerical linear algebra, Krylov methods naturally amplify eigencomponents that contribute most significantly to the finite-dimensional action of an operator, leading to rapid convergence toward invariant subspaces associated with dynamically relevant modes. Consequently, the selected eigenvectors are not merely those associated with extremal eigenvalues but those that exert the greatest influence on the operator’s behavior over the subspace generated by repeated matrix-vector products. This criterion provides a data-adaptive notion of importance that complements classical spectral ordering.

For the graph multistep diffusion operator the resulting spectrum preferentially enhances eigenvalues close to unity. Each eigenmode is weighted according to its persistence under repeated random-walk propagation. Eigenvalues satisfying |*λ*_*i*_ | ≈ 1receive substantially greater amplification than rapidly decaying modes, causing the corresponding eigenvectors to encode slowly mixing diffusion processes and persistent structural patterns. Spectrally isolated eigenvalues further improve this representation by producing well-separated invariant subspaces that are numerically stable and less susceptible to perturbations or mixing with neighboring eigenmodes. As a result, the associated embeddings capture coherent global organization rather than transient local fluctuations.

This perspective is particularly advantageous for graph representation learning because the embedding basis is determined by the effective action of the diffusion operator rather than solely by asymptotic spectral ordering. In many real-world graphs, eigenvalues often appear in dense clusters, and the associated eigenvectors may exhibit considerable sensitivity to perturbations or fail to distinguish structurally meaningful components. Selecting spectrally isolated dominant eigenpairs alleviates this ambiguity by identifying invariant directions that are both numerically robust and dynamically significant. Consequently, the resulting embedding coordinates provide a stable representation of communities, bottlenecks, and other slowly evolving structures governing information flow on the graph.

Unlike conventional spectral embedding techniques, which explicitly rank eigenvectors according to eigenvalue magnitude or Laplacian ordering, the present formulation identifies eigenpairs based on their dominant contribution to the operator’s finite-dimensional behavior. Although Krylov methods have long been employed as computational tools for large-scale eigendecomposition, their convergence properties have generally not been used as the primary criterion for selecting embedding coordinates. Existing graph embedding methods instead rely on eigendecompositions or singular value decompositions of graph-derived matrices, with the embedding dimension determined by spectral ordering. The present approach therefore introduces an alternative selection principle in which the embedding basis consists of spectrally isolated eigenpairs that are simultaneously persistent under diffusion and dominant in the Krylov-generated invariant subspace. This provides a mathematically justified representation that is closely aligned with the effective dynamics of the graph and may better preserve structurally significant information than embeddings derived solely from extremal spectral ordering.

## Code availability

The code used to perform MOSAIC embedding and analysis are available in the repository: https://github.com/TaylorResearchLab/MOSAIC.

## Acknowledgements

Research reported in this publication was supported by the NIH under award number **4U24OD038422**. Authors would also like to acknowledge Varduhi Petrosyan, Matthew Roth, Adam Resnick, Dave D. Hill, Elizabeth Goldmuntz, Kristina Cole, Sarah Tasian, Aleksandar Milosavljevic.

## Algorithm 1

Krylo-Ritz method with convergence-based ranking algorithm

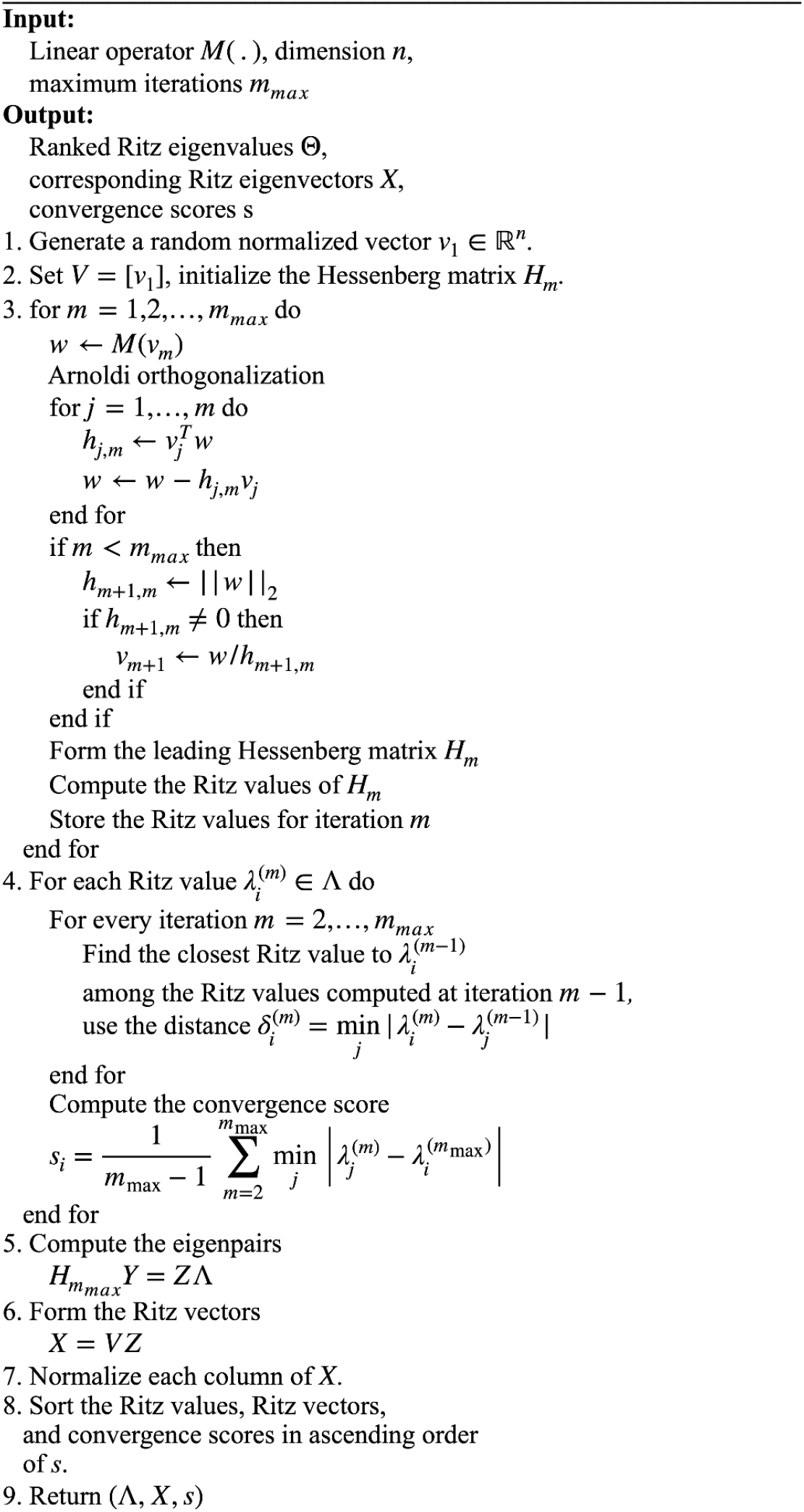

## Algorithm 2

MOSAIC algorithms

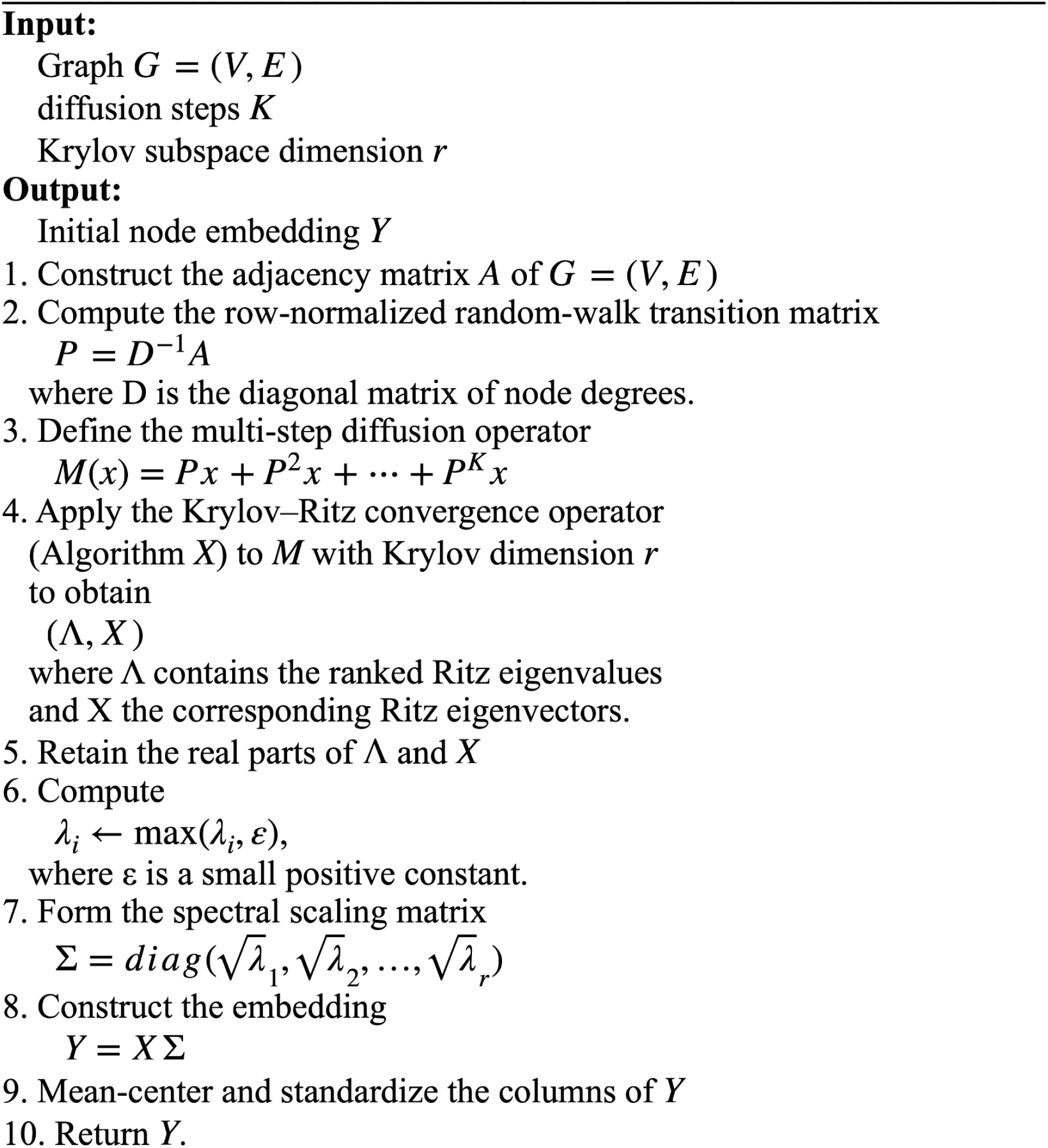

